# Interspecies malate-pyruvate shuttle drives amino acid exchange in organohalide-respiring microbial communities

**DOI:** 10.1101/379438

**Authors:** Po-Hsiang Wang, Kevin Correia, Han-Chen Ho, Naveen Venayak, Kayla Nemr, Robert Flick, Radhakrishnan Mahadevan, Elizabeth A. Edwards

## Abstract

Most microorganisms in the biosphere live in communities and develop coordinated metabolisms via trading metabolites. In this study, we sought to deconstruct the metabolic interdependency in organohalide-respiring microbial communities enriched with *Dehalobacter restrictus* (*Dhb*), using a complementary approach of computational metabolic modeling and experimental validation. *Dhb* possesses a complete set of genes for amino acid biosynthesis yet requires amino acid supplementation. We reconciled this discrepancy using Flux Balance Analysis with consideration for cofactor availability, enzyme promiscuity, and shared protein expression patterns of several *Dhb* strains. Experimentally, ^13^C incorporation assays, growth assays, and metabolite analysis of strain PER-K23 cultures were performed to validate the model predictions. The model resolved that *Dhb*’s amino acid dependency results from restricted NADPH regeneration and diagnosed that malate supplementation can replenish intracellular NADPH using malic enzyme. Interestingly, we observed unexpected export of glutamate and pyruvate in parallel to malate consumption in the strain PER-K23 cultures. Further experiments on *Dhb*-enriched consortium ACT-3 suggested an interspecies malate-pyruvate shuttle between *Dhb* and a glutamate-auxotrophic *Bacteroides* sp., reminiscent of the mitochondrial malate shunt pathway in eukaryotic cells. Altogether, this study reveals that redox constraints and metabolic complementarity are important driving forces for amino acid exchange in anaerobic microbial communities.

## Introduction

Prokaryotic microorganisms ubiquitously inhabit the biosphere, forming close associations with one another. The study of microbial communities is gaining importance due to their essential contributions in global element cycling, agriculture, bioremediation, human health and industrial biotechnology (Dolfing, 2013, Embree *et al*., 2015, Mee *et al*., 2014, Zhao *et al*., 2014). The interactions among microorganisms and their surroundings form phenotypes that can be observed in ecosystems, and are classified into three main categories: syntrophy, cross-feeding, and competition (Seth and Taga, 2014). Several methods have been proposed to elucidate these complex interactions, including microbial co-association network analysis as a function of time and other external factors (Cardona *et al*., 2016).

Metabolic complementarity is a driving force for microbial mutualism (Mori *et al*., 2016, Wintermute and Silver, 2010). While mutualism confers robustness to a microbial community, a trade-off is that the coevolved microbes within are susceptible to the loss of non-essential functions via genome streamlining, which can lead to auxotrophy in other environments (McCutcheon and Moran, 2012). As a result, isolation of microorganisms from their syntrophic partners or natural niches is often challenging, as demonstrated by the scarcity of culturable isolates in the laboratory (~2%) (Wade, 2002). An alternative approach is metabolic modeling based on microbial genomes (Manor *et al*., 2014, Roling and van Bodegom, 2014, Tan *et al*., 2015). Genome-scale constraint-based metabolic models have been increasingly used to elucidate metabolic networks at the community level (Magnúsdóttir and Thiele, 2018, Zhuang *et al*., 2011). Nevertheless, physiological information and genome annotation verification for organisms in complex communities are often lacking, which results in the inclusion of mis-annotated genes and non-gene associated reactions (Suthers *et al*., 2009), rendering simulations to be physiologically irrelevant. Integration of laboratory experiments with metabolic modeling can significantly improve the accuracy of prediction (Amador-Noguez *et al*., 2010), as demonstrated in the study of amino acid cross-feedings in synthetic *E. coli* communities (Mee *et al*., 2014).

Anaerobic organic mineralization requires tightly coupled metabolic coordination between microbes due to redox and thermodynamic constraints (McInerney *et al*., 2009, Sieber *et al*., 2012). The presence of external electron acceptors enables more complex communities to develop. Organohalide-respiring microbial communities are great models to study metabolic interdependency between microbes because they are often co-inhabited by acetogens, fermenting bacteria, methanogens, sulfate-reducing bacteria, and organohalide-respiring bacteria (OHRB) (Adrian and Loeffler, 2016, Duhamel and Edwards, 2007). *Dehalobacter restrictus* (*Dhb*) strains are specialized in respiring a variety of organohalides (Holliger *et al*., 1998, Justicia-Leon *et al*., 2012, Tang *et al*., 2016, van Doesburg *et al*., 2005, Wang *et al*., 2014, Wong *et al*., 2016, Yoshida *et al*., 2009). All isolated *Dhb* strains, with the exception of strain UNSWDHB, require the addition of either amino acids or parent culture supernatants to support growth (Holliger *et al*., 1998, Wang *et al*., 2016), indicating unexplored nutrient cross-feedings from symbiotic partners in natural habitats, e.g. the *Bacteroides* sp. (*Bac*) in the strain CF-enriched consortium ACT-3 (Tang and Edwards, 2013b). However, comparative genomic analysis and refined metabolic annotation suggested that *Dhb* possesses a complete set of genes to synthesize all amino acids, including a salvage pathway to obtain serine from threonine (Tang *et al*., 2016, Wang *et al*., 2016).

In this study, we first explored the metabolic dependency of *Dhb* using its genome-scale metabolic model (Correia *et al*., 2018), which was built based on a highly annotated and experimentally refined *Dhb* genome (Wang et al, 2016), along with shared expression patterns in proteomic datasets of *Dhb* strains PER-K23, UNSWDHB, DCA and CF (Jugder *et al*., 2016, Rupakula *et al*., 2013, Rupakula *et al*., 2015, Tang and Edwards, 2013a). The model simulations consider cofactor availability, enzyme promiscuity, physiological redox conditions, and further model constraints based on experimental values obtained in this study. We resolved that *Dhb*’s amino acid dependency results from restricted NADPH regeneration, which can be restored with malate supplementation via the function of NADP^+^-dependent malic enzyme. The strain PER-K23 cultures grown on the model-resolved medium exhibited an unexpected export of pyruvate, glutamate, and other amino acids in parallel to malate consumption. Further experimental analysis on the strain CF-enriched consortium ACT-3 revealed that *Dhb*’s specialized malate requirement is likely a consequence of genome streamlining, which was driven by co-adaption with a glutamate-auxotrophic, malate-producing *Bacteroides* sp. to effectively accomplish intercellular trading of reducing equivalents (i.e. NADH and NADPH) via organic acid exchange.

## Materials and Methods

All Chemicals were ordered from Sigma-Aldrich (Oakville, ON, Canada) at highest purity available unless specified otherwise.

### Flux balance analysis (FBA)

The experimentally refined genome of *D. restrictus* strain CF (accession no. NC_018866) (Wang et al, 2016) was used to reconstruct a draft *Dhb* genome-scale metabolic model (Correia *et al*., 2018). Flux balance analysis and flux variability analysis simulations were conducted with COBRApy (Ebrahim *et al*., 2013). Constraints were applied to the genome-scale metabolic model based on five considerations to improve the accuracy of flux distributions with *Dhb*: (i) consideration of cofactor availability for the corresponding metabolic reactions; (ii) shared features in available proteomes of *Dhb* strains (**Table S1**); (iii) consideration of cellular redox state under given growth conditions; (iv) consideration of potential promiscuous enzyme activity to rescue missing pathways; (v) integration of experimental values from culture growth assays and metabolite profile analysis. Based on these considerations, constraints were applied to the model by the following four actions: (a) inactivating targeted reactions, (b) limiting the direction of targeted reactions (making reactions irreversible), (c) applying experimentally relevant metabolite flux, and (d) disabling/enabling the export of metabolites (**Table S2**).

### Microbial cultures and growth conditions

*Dhb* strain PER-K23 was provided by the Löffler Lab at University of Tennessee (Knoxville, USA). *Escherichia coli* strain BL21(DE3) was purchased from New England Biolabs Ltd. Consortium ACT-3 was originally enriched from 1,1,1-trichloroethane-contaminated groundwater in 2001 from a northeastern United States industrial area (Grostern and Edwards, 2006), and a subculture (1.8 l) was adapted to respire chloroform (Grostern *et al*., 2010). *E. coli* strain BL21 was grown on LB broth except in the ^13^C incorporation assay. The strain PER-K23 cultures and the consortium ACT-3 are maintained in a FeS-reduced, bicarbonate-based mineral medium described previously (Grostern *et al*., 2010, Wang *et al*., 2016). The growth assays for the strain PER-K23 cultures and consortium ACT-3 were performed following the established protocol reported previously with some modifications (Tang and Edwards, 2013a, Wang *et al*., 2016) (described in **SI**).

### ^13^C incorporation assay

The ^13^C incorporation assay was performed following a previous study on amino acid biosynthesis of *Dehalococcoides* with some modifications (Zhuang *et al*., 2014) (described in **SI**).

### DNA extraction, quantitative PCR, and Illumina amplicon sequencing

Culture DNA was extracted from 2 ml samples. Cells were harvested by centrifugation at 16 000 x g for 10 min at 4°C. Since *Dhb* cell pellets are easily resuspended, in each tube, most of supernatant was remove (1.9 ml), and the cell pellets were resuspended using the remaining supernatant (0.1 ml), and the DNA was extracted using the MO BIO PowerSoil^®^ DNA isolation kit following the manufacturer’s recommendations. Real-time quantitative polymerase chain reaction (qPCR) assays were performed to track the gene copy numbers of *Dhb* using specific 16S rRNA gene primers reported previously (F: 5’-GAT TGA CGG TAC CTA ACG AGG-3’; R: 5’-TAC AGT TTC CAA TGC TTT ACG-3’) (Puentes Jácome and Edwards, 2017) (described in **SI**). For the 16S rRNA gene-based population analysis, the ACT-3 sub-transfer cultures (2 ml) from each triplicate trials were combined, and the DNA were extracted as described above. The DNA samples were sent to the Genome Quebec Innovation Centre (McGill University, Canada) for Illumina MiSeq amplicon sequencing. After sequencing, the raw data were processed following an established pipeline described previously (Chen *et al*., 2018). The assemblage of pair-end reads, primer removal, quality filtering, chimera and singleton detections, and read number normalization were implemented using the sequence analysis tool USEARCH, and the taxonomic assignment of OTUs was performed against the Silva database (version 128; https://www.arb-silva.de/documentation/release-128/). The taxonomic assignment and abundance of individual OTUs is available in **Table S3**.

### Phylogenetic analysis

The species tree was created with a modified method recently published (Hug *et al*., 2016). Briefly, 10 ribosome protein subunits in the selected bacterial species were independently aligned with MAFFT v7.245, trimmed to remove unaligned N and C termini residues using default parameters with Gblocks version 0.91b (Talavera and Castresana, 2007), and concatenated to reconstruct a maximum likelihood tree with 100 bootstrap values using PHYML v3.2.0 (Guindon *et al*., 2010). Ortholog groups were predicted via OrthoMCL (Fischer *et al*., 2011) and OrthoDB (Zdobnov *et al*., 2016). Enzyme orthologs were mapped to the species tree with Evolview v2 (He *et al*., 2016).

### Enzyme activity assays

The conditions for cell extract preparation and dehalogenase activity assays were described previously (Wang et al, 2016), and o-phosphoserine phosphatase activity assay were conducted following established protocols with some modifications (described in **SI**) (Kuznetsova *et al*., 2006). The enzyme activity is defined as μmol product produced min^−1^ mg protein^−1^.

### Analytical procedures

Chlorinated hydrocarbons were measured by injecting a 0.3 ml headspace sample into a Hewlett-Packard 5890 Series II GC fitted with a GSQ column (30-m-by-0.53-mm [inner diameter] PLOT column; J&W Scientific, Folsom, CA) as described previously (Wang et al, 2016). For the metabolite profile analysis, in an anaerobic chamber (Coy), each culture was sampled (0.2 ml), and gently filtered through a 0.1 μm-pore-size syringe filter (Millipore). The flow-through was collected in a plastic microcentrifuge tube, followed by a centrifugation at 16 000 x g for 10 min at 4°C. The supernatants were stored at −80°C before analysis. The amount of acetate, malate and pyruvate from *Dhb* strain PER-K23 cultures was determined by HPLC using an ICS5000 system (Thermo scientific) equipped with an Aminex HPX-87H column (BioRad) connected to a UV detector. Each sample (25 μl) was injected onto the column incubated at 35°C, using 5 mM H_2_SO_4_ eluent at a flow rate of 0.6 ml^−1^ min with the UV wavelength set to 210 nm. Amino acids and organic acids were detected using Liquid chromatography Electrospray-coupled high resolution mass spectrometry (LC-ESI-HRMS) with a Dionex UHPLC system and a Q-Exactive mass spectrometer (Thermo Scientific) equipped with a HESI II source (Thermo Scientific) and a Micro-splitter valve (IDEX Health & Science) (described in **SI**).

## Results and Discussion

### Serine biosynthesis in *Dhb* via threonine

Our research efforts have been devoted to refining the genome annotation on the central metabolism and biosynthesis of amino acids/cofactors in *Dhb* (Wang et al, 2016). Common features among *Dhb* genomes are gaps in the TCA cycle and serine biosynthesis. To increase accuracy in model reconstruction, we analyzed the presence/absence of enzyme orthologs involved in these metabolic pathways in a range of OHRB species, *Firmicutes* species, *Bacteroides* species, and selected model organisms in Bacteria (**Figure 1**). Malate dehydrogenase and succinate dehydrogenase/fumarate reductase genes appear to have been recently lost in *Dhb*. The membrane-bound transhydrogenase PntAB is absent in *Dhb* and many Firmicutes species, reducing their flexibility in NADH/NADPH metabolism. Interestingly, the orthologs of o-phosphoserine phosphatase (SerB; EC 3.1.3.3), the enzyme catalyzing the final step in the classical serine biosynthesis pathway (Greenberg and Ichihara, 1957), are notably absent in Firmicutes. Conservation of SerA in most Firmicutes likely results from the overlapping biosynthesis pathway between serine and pyridoxine, an essential cofactor involved in central metabolism (**Figure 1**) (Lam and Winkler, 1990). Previously, we found a promiscuous serine hydroxymethyltransferase (Shmt; EC 2.1.2.1) possessing threonine aldolase activity in *Dhb*, which allows serine salvage from threonine (**Figure 2A**). Moreover, this promiscuous serine hydroxymethyltransferase is expressed in all available *Dhb* proteomes (**Table S1**). To exclude the possibility that a phylogenetically distant SerB or a promiscuous phosphatase with SerB-like activity is present in *Dhb*, we first examined the growth of strain PER-K23 cultures with only acetate or with acetate and serine. Dechlorination was only observed in cultures supplemented with both acetate and serine (**Figure S1A**). We then examined potential promiscuous phosphatase activity in *Dhb* cell lysates. While we observed dehalogenase activity in assays containing the *Dhb* cell lysates, SerB-like activity was only observed in assays containing *E. coli* cell lysates (positive control) (**Figure 2B**). These preliminary results indicate that SerB-dependent classical serine biosynthesis pathway is not functioning in *Dhb* cells.

**Figure 1.**
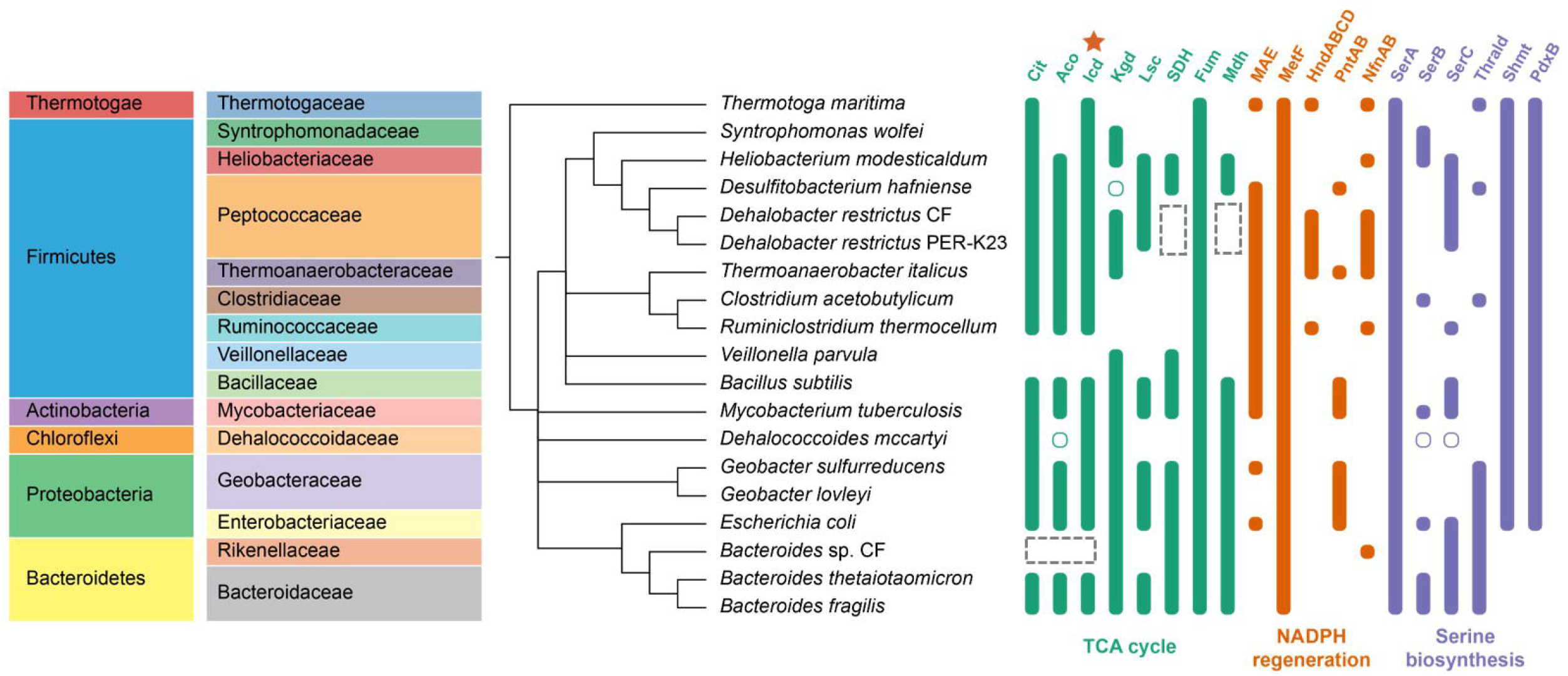
Presence/Absence of enzyme orthologs in the TCA cycle, NADPH regeneration, and serine biosynthesis mapped to a species tree of organohalide-respiring bacteria and selected organisms. Open circles represent the enzyme activity are present in the corresponding organisms based on experimental evidences but without annotated enzyme orthologs; the dashed line boxes highlight the missing TCA cycle enzymes in *Dehalobacter* strains and *Bacteroides* sp. CF. Note that isocitrate dehydrogenase (Icd) in the TCA cycle is also involved in NADPH regeneration (orange star). Abbreviations: SDH, succinate dehydrogenase/fumarate reductase; Mdh, malate dehydrogenase, Cit, citrate synthase; Aco, aconitase; MAE, malic enzyme; MetF, 5,10-methelene-tetrahydrofolate reductase/dehydrogenase; HndABCD (or HymABC), NADP^+^-reducing hydrogenase; NfnAB, NADP^+^:ferredoxin oxidoreductase; PntAB, membrane-bound transhydrogenase; SerA, 3-phosphoglycerate dehydrogenase; SerB, o-phosphoserine phosphatase; SerC, o-phosphoserine aminotransferase; Thrald, threonine aldolase; Shmt, serine hydroxymethyltransferase; PdxB, erythrose-4-phosphate dehydrogenase.

**Figure 2.**
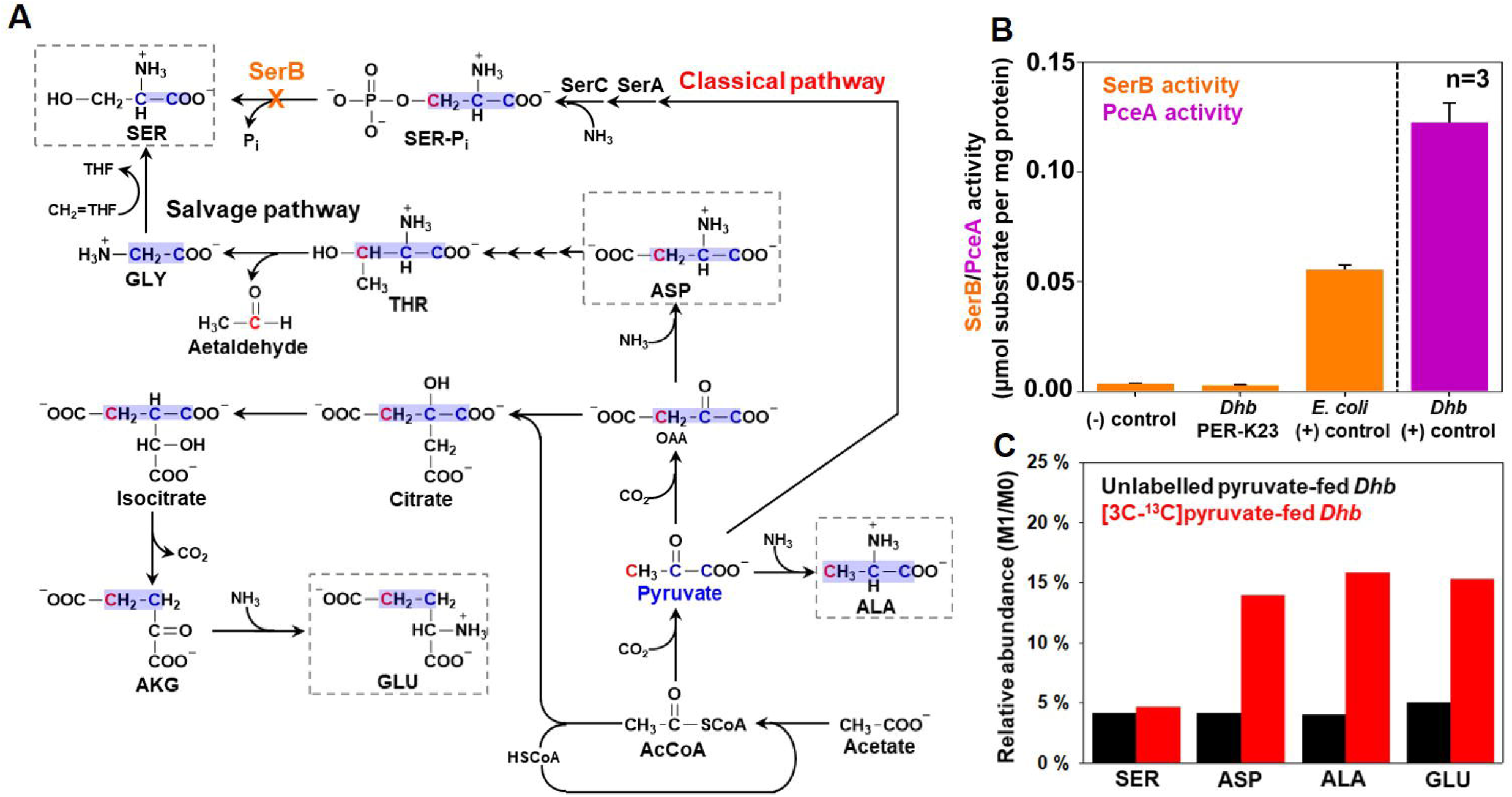
Serine biosynthesis in *Dehalobacter restrictus* via threonine. **(A)** Schematic of amino acid biosynthesis (alanine, aspartate, glutamate, and serine; in dashed boxes) and carbon incorporation in *Dehalobacter restrictus* (*Dhb*) using [3-^13^C]pyruvate as the precursor. Serine is synthesized via the salvage pathway in *Dhb* due to the lack of SerB (highlighted in orange), the enzyme catalyzing the final step in the classical serine biosynthesis pathway. The ^13^C-labelled carbon originating from [3-^13^C]pyruvate is shown in red and the carbons derived from pyruvate is highlighted in cyan. **(B)** SerB activities in cell lysates of strain PER-K23 and *E. coli*, respectively. (−) control, cell-lysate-free control. Tetrachloroethene dehalogenase (PceA) activity was used as a quality control of strain PER-K23 cell lysates. Data are means ± SE of three replicates in each experiment as shown on figures. **(C)** Relative abundance of different mass isotopomers (M1/M0) of serine, aspartate, alanine, and glutamate obtained from strain PER-K23 cells cultivated on the defined medium supplemented with unlabeled pyruvate or [3-^13^C]pyruvate. Abbreviations: AcCoA, acetyl-CoA; AKG, α-ketoglutarate; CH_2_=THF, 5,10-methylenetetrahydrofolate; MAL, OAA, oxaloacetate; SER-P_i_, o-phosphoserine; SerA, phosphoglycerate dehydrogenase; SerB, phosphoserine phosphatase; SerC, phosphoserine aminotransferase.

Subsequently, we followed ^13^C incorporation to elucidate serine biosynthesis in *Dhb*. [3-^13^C]pyruvate or unlabeled pyruvate (5 mM), a precursor of serine in the classical pathway (**Figure 2A**) (Grundy and Henkin, 2002), was supplemented to Holliger medium (Holliger *et al*., 1998, Wang *et al*., 2016) containing 1 mM acetate and 0.1 mM of arginine, histidine, and threonine to cultivate strain PER-K23 cultures. Therefore, the 3-^13^C on pyruvate would be incorporated into serine if *Dhb* synthesizes serine via the classical pathway (**Figure 2A**), and MS analysis of isotopomer distribution would reveal an enrichment in relative abundance of the M1 isotopomer of serine (M1/M0). By contrast, the M1 serine isotopomer would remain at natural abundance if *Dhb* salvages serine from threonine. Consistently, serine obtained from both unlabeled pyruvate-fed and [3-^13^C]pyruvate-fed *Dhb* cells revealed a comparable M1/M0 (4.2~4.7%) (**Figure 2C**). By contrast, alanine, aspartate, and glutamate obtained from [3-^13^C]pyruvate-fed *Dhb* cells revealed at least 3-fold enrichment in M1/M0 (~15%). Altogether, based on genome annotation and multiple lines of experimental evidence, the classical serine biosynthesis pathway is absent in *Dhb*, and serine is synthesized from the salvage pathway. Interestingly, this salvage pathway is likely a major route for serine biosynthesis in SerB-lacking Firmicutes and in *Geobacter* spp. (**Figure 1**) (Sung, et al 2006).

### Restricted NADPH regeneration in *Dhb*

The absence of the classical serine biosynthesis pathway in *Dhb* can result in many metabolic defects because serine is the cellular C_1_ pool donor and the precursor of purines and other amino acids (**Figure 3A**) (Fan *et al*., 2014). Recent studies also discovered that serine can support NADPH regeneration via the folate cycle (**Figure 3A**) (Fan *et al*., 2014, Tedeschi *et al*., 2013). Furthermore, biosynthesis of serine and serine-derived amino acids through the classical pathway is NADPH-independent (embedded table in **Figure 3B**) (Grundy and Henkin, 2002). By contrast, using threonine to synthesize serine would elevate cellular demand of NADPH for amino acid synthesis by approximately 30%, assuming a protein composition similar to that of *Bacillus subtilis* (Dauner and Sauer, 2001).

**Figure 3.**
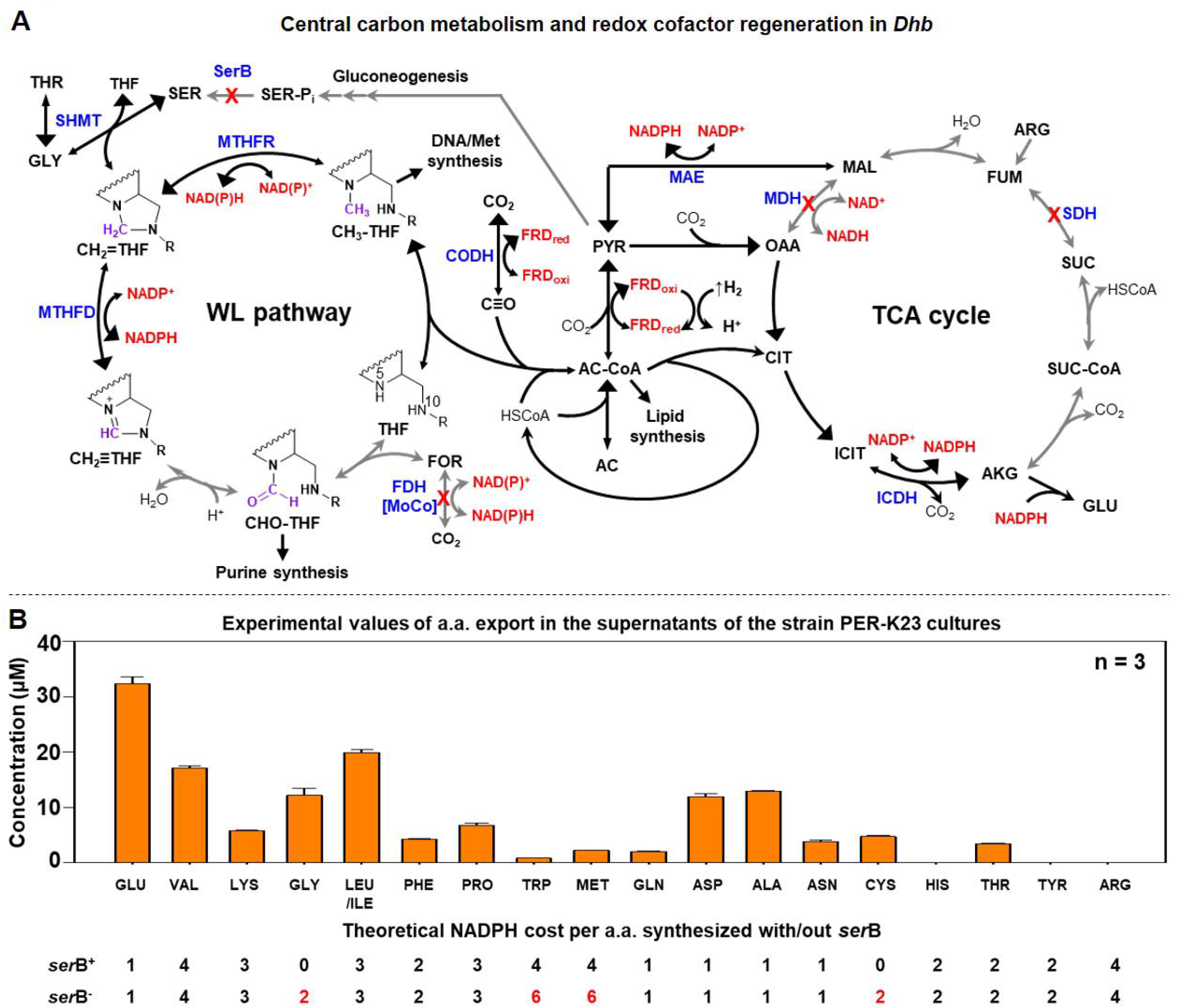
Amino acid dependency of *Dehalobacter restrictus* results from restricted redox metabolism (NADPH and ferredoxin). **(A)** Proposed central carbon metabolism and redox cofactor regeneration system in *Dehalobacter restrictus* (*Dhb*) with acetate and malate as the carbon sources. The name of enzymes and the missing cofactors (in square bracket) are shown in blue. The red crosses (X) represent the missing genes or missing cofactors in the *Dhb* genome. NAD(P)^+^/NAD(P)H are shown in red beside the corresponding metabolic reactions. The black arrows represent the metabolic reactions involved in NADPH and ferredoxin regeneration, while grey arrows represent the reactions not involved. The double-headed arrows indicate reversible reactions, and the bigger arrowheads represent the direction of reactions under physiological conditions. **(B)** Amino acid profile in the supernatants of strain PER-K23 cultures after the consumption of 5 mM TCE (**Figure 4A**). The amounts of NADPH required to synthesize each amino acid via the classical pathway (SerB^+^) or salvage pathway (SerB^−^) are listed below. Data are means ± SE of three replicates in each experiment as shown on figures. Abbreviations: AC, acetate; AC-CoA, acetyl-CoA; AKG, α-ketoglutarate; CH2=THF, 5,10-methylenetetrahydrofolate; CH2=THF, 5,10-methenyltetrahydrofolate; CH3-THF, 5-methyltetrahydrofolate; CHO-THF, 5-formyl-tetrahydrofolate; CIT, citrate; FOR, formate; FUM, fumarate; FRD, ferredoxin; ICIT, isocitrate; MAL, malate; OAA, oxaloacetate; PYR, pyruvate; SER-P_i_, O-phosphoserine; SUC, succinate; SUC-CoA, succinyl-CoA; THF, tetrahydrofolate. Abbreviations for enzymes: CODH, carbon monoxide dehydrogenase; FDH, formate dehydrogenase; FUM/SDH, fumarate reductase/succinate dehydrogenase; MAE, malic enzyme; MTHFD, 5,10-methylenetetrahydrofolate dehydrogenase; MTHFR, 5,10-methylenetetrahydrofolate reductase; MDH, malate dehydrogenase; SerB, o-phosphoserine phosphatase; WL pathway, Wood–Ljungdahl pathway (or folate cycle).

According to *Dhb* genome annotation, seven potential enzyme reactions can contribute to NADPH regeneration (**Figure 3A**), including a putative ferredoxin:NADP^+^ oxidoreductase (NfnAB; EC 1.6.1.4); a putative NADP^+^-reducing hydrogenase (HndABCD; EC 1.12.1.3); isocitrate dehydrogenase (EC 1.1.1.42); NADP-dependent malic enzyme (MAE; EC 1.1.1.40); MoCo-dependent NADP^+^-specific formate dehydrogenase (FdhAB; EC 1.2.1.43); and 5,10-methylenetetrahydrofolate reductase/dehydrogenase (MetF; EC 1.5.1.20; FolD; EC 1.5.1.5). However, *Dhb* cannot synthesize the molybdopterin cofactor (Wang *et al*., 2016) to functionalize formate dehydrogenase. Also, given that *Dhb* employs the TCA cycle and Wood–Ljungdahl pathway for anabolism but not respiration, the generated NADPH from isocitrate dehydrogenase and 5,10-methylenetetrahydrofolate reductase/dehydrogenase is insufficient to support *Dhb* anabolism using simple carbon sources like acetate. Finally, the putative NADP^+^-reducing hydrogenase was not expressed in all the available *Dhb* proteomes, with the putative ferredoxin:NADP^+^ oxidoreductase only being expressed in the strain UNSWDHB proteome (**Table S1**). Consistently, strain UNSWDHB was reported to grow on acetate as the sole carbon source (Wong *et al*., 2016) and was isolated from an acetate/H_2_-fed enrichment, while strains PER-K23, CF, and DCA were isolated from lactate-fed enrichment cultures (Grostern and Edwards, 2006, Holliger *et al*., 1993). Such difference in ferredoxin:NADP^+^ oxidoreductase expression among *Dhb* strains is likely a result of niche specialization driven by bacterial epigenetic change (e.g. DNA methylation) responding to environmental conditions (Casadesús and Low, 2006).

Among the seven NADPH-regenerating enzymes, MAE can replenish the intracellular NADPH pool with malate supplementation (**Figure 3A**). Also, *Dhb* genomes possess malate permease (*maeP*; Accession K4LDY2) to uptake malate, with MAE being significantly expressed in the proteomes of all *Dhb* strains (**Table S1**). Thus we examined if malate supplementation can support *Dhb* growth on the acetate-based defined medium. Consistently, strain PER-K23 cultures supplemented with acetate and malate showed a three-fold faster dechlorination rate than that of cultures supplemented with acetate and serine (**Figure S1A**). Supplementing acetate, malate and serine to strain PER-K23 cultures resulted in the highest dechlorination rate (100 μM bottle^−1^ day^−1^). Therefore, the lack of SerB and restricted NADPH regeneration system is likely the result of *Dhb*’s amino acid dependency.

### Unexpected pyruvate export in *Dhb* cultures

To examine the hypothesis of restricted NADPH regeneration in *Dhb*, we performed three consecutive 1% transfers of strain PER-K23 cultures using the defined medium supplemented with both malate and serine but not acetate. Strain PER-K23 cultures were able to deplete 1 mM TCE after three 1% transfers (**Figure S1B**). We then examined long-term growth of strain PER-K23 cultures. The cultures readily depleted three feedings of 1 mM TCE (**Figure 4A**). qPCR analysis revealed that *Dhb* cell density in the cultures increased by 100-fold after the consumption of 3 mM TCE (~2 x 10^7^ cells ml^−1^) (**Figure 4A**). The cell growth parameter (6.3 ± 0.02 x 10^12^ cells mol^−1^ Cl^−^ released) is in good agreement to that of strains CF and UNSWDHB reported previously (Grostern *et al*., 2010, Wong *et al*., 2016). However, after the fourth feeding, the cultures displayed a significant drop in dechlorination rate. Before each feed, cultures were purged with N_2_/CO_2_ to remove cis-dichloroethene, to adjust the pH, and to replenish the medium with H_2_. Moreover, the metabolite profile of culture supernatants revealed that most malate and serine (~90%) were not consumed (**Figure 4B**). Thus the lagging dechlorination is not due to either the shortage of electron donor/carbon sources, accumulation of electron acceptor, or acidic pH. Unexpectedly, the metabolite profile of strain PER-K23 culture supernatants revealed pyruvate production in parallel to malate/serine consumption, which can be produced via the function of MAE, serine deaminase (SdaA; EC 4.3.1.17), or pyruvate:ferredoxin oxidoreductase, expressed in all *Dhb* proteomes (**Figure 3A; Table S1**). This unexpected pyruvate export suggests an unfavourable sink for pyruvate in *Dhb*. Indeed, the lack of malate dehydrogenase and succinate dehydrogenase/fumarate reductase genes prevents *Dhb* to ferment pyruvate to succinate, and the absence of the classical serine pathway prevents pyruvate to enter the Wood–Ljungdahl pathway via serine (**Figure 3A**). Furthermore, from a thermodynamics perspective, MAE-mediated malate decarboxylation is unfavorable under standard conditions (**Table 1**). Therefore, pyruvate (reaction product) must be exported to drive the malate-dependent NADPH regeneration.

**Figure 4.**
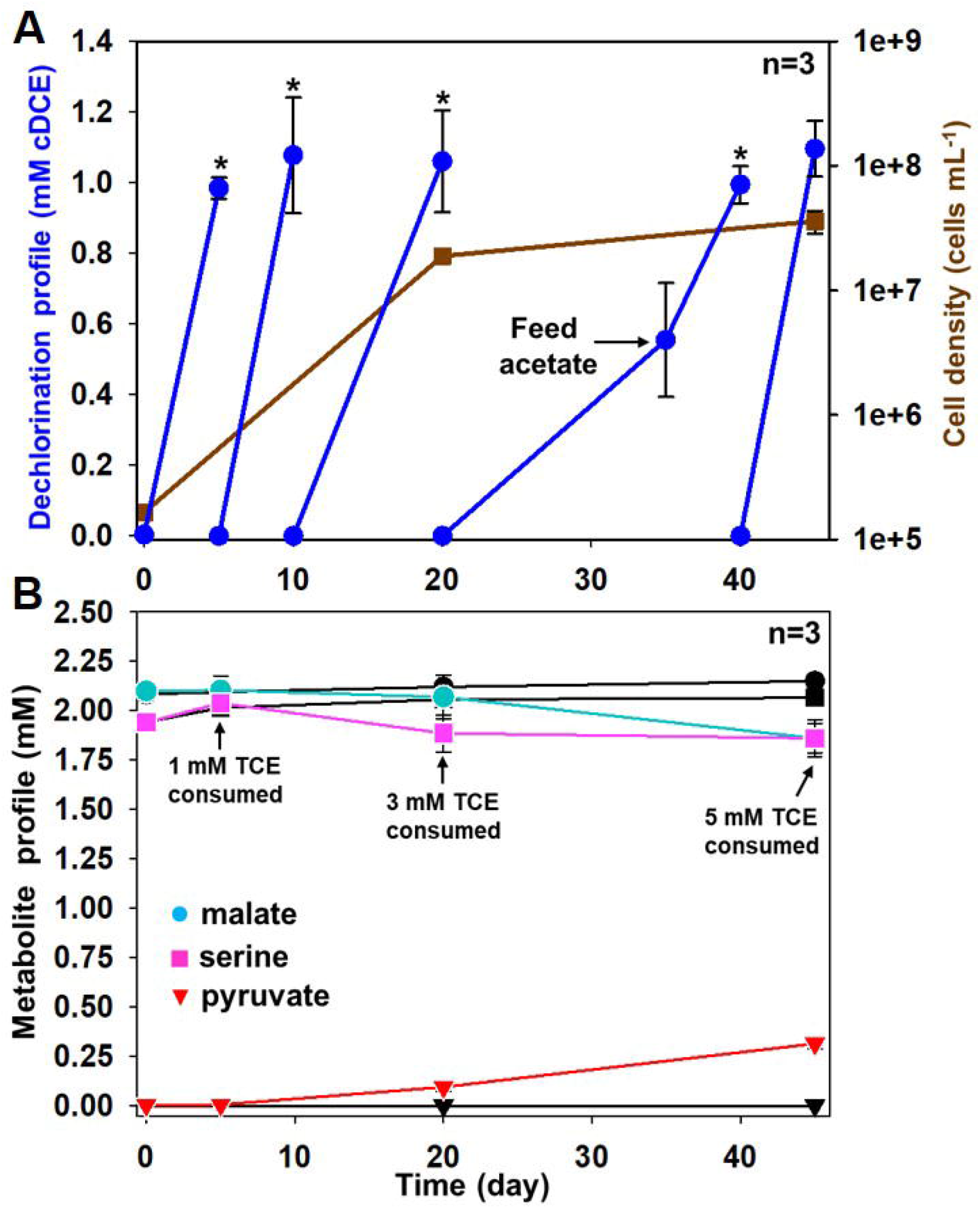
Growth assays of *Dehalobacter restrictus* strain PER-K23. **(A)** Cumulative tetrachloroethene (TCE) dechlorination profile (blue circle) and cell growth (brown square) by the strain PER-K23 cultures cultivated on the defined medium amended with malate and serine. When the fed TCE is depleted (*), the cultures were purged with 80% N_2_/CO_2_ to remove dechlorination product cis-dichloroethene (cDCE) and re-fed with H_2_ and TCE (1 mM). **(B)** Time-course metabolite profile (malate, circle; serine, square; pyruvate, triangle) of the culture supernatants. The black-colored plots represent the killed controls. The slightly gradual increase in substrate concentration in the killed controls is due to water evaporation during culture purging. The cumulative TCE consumption is also shown. Data are means ± SE of three replicates in each experiment as shown on figures.

**Table 1.** Overall reaction and eQuilibrator-estimated Δ_*r*_G’^o^ mentioned in this study.

|  | **Reaction** | **$\Delta_r G'^0$**<br>**(kJ/mol)** |
| --- | --- | --- |
| **LAC fermentation by *Bacteroides*** | Lactate + 0.5 H<sub>2</sub>O → 0.5 Acetate + 0.5 Malate + H<sub>2</sub> | 23 |
| **Chloroform dechlorination by *Dehalobacter*** | CF + H<sub>2</sub> → DCM + HCl | -173 |
| **Malic enzyme reaction** | Malate + NADP<sup>+</sup> → Pyruvate + CO<sub>2</sub> + NADPH | 14 |
| **syntrophic LAC fermentation by *Bacteroides* and *Dehalobacter*** | Lactate + 0.5 H<sub>2</sub>O + CF → 0.5 Acetate + 0.5 Malate + DCM + HCl | -150 |
| **Interspecies NADH/NADPH exchange achieved by the malate-pyruvate shuttle** | NADH + NADP<sup>+</sup> → NAD<sup>+</sup> + NADPH | -2.3 |
| **AcCoA-dependent ATP synthesis (substrate-level phosphorylation)** | AcCoA + ADP + Pi → Acetate + ATP + CoA | -3.6 |

### Deconstructing *Dhb* metabolism using complementary FBA and experimental validation

We then simulated *Dhb* metabolism using the model constrained by experimental values for cell growth and substrate consumption (H_2_, TCE, malate, and serine) in strain PER-K23 cultures at Day 20 (**Figure 4B**). In addition to pyruvate, the resulting model also predicted significant CO export (**Figure S2A**). Since we purged the cultures before each feed, CO would not accumulate to cause cell toxicity. Instead, CO production via ferredoxin-dependent CO dehydrogenase (CooS; EC 1.2.7.4) indicates excessive reduced ferredoxin (FRD_red_) accumulation (**Figure 3A**). Given that *Dhb* grows under conditions with excessive H_2_, the model predicted excessive FRD_red_ production from the FRD-reducing hydrogenases (Ech and HymABC) expressed in all available *Dhb* proteomes (**Tables S1, S2**). FRD_red_ accumulation will inhibit conversion of pyruvate to acetyl-CoA in central carbon metabolism while enhancing the reverse reaction (**Figure 3A**). Since acetyl-CoA is the substrate for citrate synthase in the oxidative TCA cycle, acetyl-CoA shortage would disable citrate synthesis, preventing pyruvate from entering the TCA cycle for amino acid biosynthesis. Therefore, the model predicted CO export to consume excessive FRD_red_, enabling acetyl-CoA synthesis via pyruvate. Accordingly, acetate supplementation to strain PER-K23 cultures would replenish the acetyl-CoA pool in *Dhb* and ameliorate the imbalanced redox potential. Consistently, acetate supplementation (1 mM) at Day 35 restored the lagging dechlorination, and strain PER-K23 cultures were able to deplete the remaining TCE and a subsequent feeding in 10 days (**Figure 4A**), along with a significant pyruvate production (0.3 mM) in parallel to the consumption of acetate (0.15 mM), malate (0.25 mM) and serine (0.1 mM). In the first three feeding cycles, strain PER-K23 cells likely used the residual acetate in the medium components (≤ 0.1 mM based on HPLC analysis) to synthesize acetyl-CoA. Therefore, the limited sink for pyruvate and excessive FRD_red_ production in *Dhb* resulted in pyruvate export.

After resolving FRD metabolism in *Dhb*., we further constrained the model with experimental values of strain PER-K23 cultures at Day 45, including the pyruvate export (**Figure 4B**). However, the resulting model revealed excessive NADPH production from the consumed malate (0.25 mM) via the MAE reaction, unless the model allowed (a) export of citrate in the oxidative TCA cycle to prevent NADPH production via isocitrate dehydrogenase (**Figure 3A**) or (b) export of amino acids to consume NADPH (e.g. glutamate, valine, leucine, proline) (**Figure S2B**). Congruently, metabolite profiling of the culture supernatants at Day 45 revealed apparent production of glutamate (33 μM), valine (18 μM), and iso/leucine (20 μM) against the killed control (**Figure 3B**), and these amino acids were not detected (< 0.1 μM) in the culture supernatants of *E. coli* cultures cultivated on the defined medium at the same cell density (4 x 10^7^ cells ml^−1^). Thus, *Dhb* appears to export amino acids to consume excessive NADPH when a NADPH-regenerating source like malate is abundant in the growth medium. Additionally, some alanine and aspartate were also detected in the culture supernatants. Since these two amino acids are downstream derivatives of pyruvate (**Figure 2A**), *Dhb* likely exports them to prevent excessive NADPH production by isocitrate dehydrogenase (**Figure 3A**). However, citrate was not detected in the culture supernatants (<0.1 μM). Altogether, our data suggested that both acetate and malate are required to sustain the minimal growth of *Dhb*, and serine supplementation can reduce the NADPH requirements in *Dhb* anabolism.

In this study, our integrated approaches have resolved the metabolic dependency of *Dhb*, which brings previously identified arginine and threonine requirements of strain PER-K23 cultures into context (Holliger *et al*., 1998). Due to the lack of SerB, *Dhb* utilizes threonine to synthesize serine and its derived amino acids. Also, due to the restricted NADPH regeneration system, arginine is degraded to malate via fumarate for NADPH regeneration (**Figure 3A**) (Cunin *et al*., 1986). By contrast, when malate is available, *Dhb* becomes an amino acid producer, exporting amino acids to consume excessive NADPH (**Figure 3B**), along with the decarboxylated product, pyruvate (**Figure 4B**). This finding has explained the common co-occurrence of the amino acid- and pyruvate-fermenting *Sedimentibacter* spp. in *Dhb*-enriched consortia (Imachi *et al*., 2016, Maphosa *et al*., 2012, Tang *et al*., 2012, Yoshida *et al*., 2009). However, amino acid export also limits *Dhb*’s biomass generation, resulting in the unusual low cell growth (**Figure 4A**). Compared to *Dhb*, other OHRB possess a more flexible system in NADPH regeneration (**Figure 1**). For example, the classical serine biosynthesis pathway is present in *Dehalococcoides* based on a ^13^C incorporation experiment, and an incomplete Wood–Ljungdahl pathway can support NADPH regeneration (**Figure 3A**) (Zhuang *et al*., 2014). *Geobacter lovleyi*, while lacking SerB, possesses a functional TCA cycle to respire acetate and generate sufficient NADPH using isocitrate dehydrogenase (**Figure 3A**) (Galushko and Schink, 2000, Sung *et al*., 2006). Finally, *Desulfitobacterium hafniense*, in contrast to its close relative *Dhb*, retains malate dehydrogenase and fumarate reductase genes in the TCA cycle, allowing pyruvate fermentation to malate for NADPH regeneration via MAE (**Figure 3A**) (Peng *et al*., 2012).

### Metabolic interdependencies in the *Dhb*-enriched consortium

After characterizing *Dhb’s* metabolic dependency, we wondered if these nutrients are available in *Dhb*-enriched microbial communities. Acetate is the final product of acetogenic and fermenting bacteria that often co-exist in organohalide-respiring consortia (Heimann *et al*., 2006), while the presence of malate or serine has not been reported before. Therefore, we analyzed the metabolites in supernatants of the strain CF-enriched consortium ACT-3. After lactate and chloroform were fed, time-course metabolite profile revealed malate accumulation (~0.5 μM) in the ACT-3 supernatants along with chloroform dechlorination to dichloromethane (**Figure S3**). The spiked malate concentration lasted until the fed lactate was depleted, with malate not being detected in the 1 M lactate stock based on LC-MS analysis (< 50 nM). Therefore, malate is a natural substrate for *Dhb* in ACT-3. The detection of malate production in ACT-3 brings the present malate permease gene in *Dhb* genomes and the consistent MAE expression in all *Dhb* strains into context. Also, finding available malate in ACT-3 suggests that *Dhb* has sufficient NADPH to drive a more NADPH-consuming pathway (SerB-independent pathway) for amino acid synthesis (the embedded table in **Figure 3B**). However, in contrast to strain PER-K23 cultures, no amino acid was detected (<0.1 μM) in ACT-3 supernatants. Trace pyruvate is present, but the concentration remained unchanged throughout dechlorination (**Figure S3**). These data suggested that other microbial populations consume the amino acids and pyruvate exported by *Dhb*.

We then managed to identify the malate producer in ACT-3. Based on 16S rRNA pyrotag sequences (accession SRX181448), *Bac* is a dominant population in the ACT-3 (Grostern and Edwards, 2006, Grostern *et al*., 2010, Tang *et al*., 2012). Congruently, fermentative malate/fumarate production by *Bacteroides* spp. has been studied extensively (Chen and Wolin, 1981, Macy *et al*., 1975, Macy *et al*., 1978, Miller, 1978). Due to the lack of the heme biosynthesis pathway, *Bacteroides* spp. only ferments glucose to acetate, H_2_, and malate/fumarate via lactate (Chen and Wolin, 1981), unless exogenous heme is present to support further malate fermentation to succinate (**Figure 5A**). Consistently, the closed genome of *Bac* (accession CP006772) reveals complete pathways for lactate fermentation to acetate and succinate (Tang and Edwards, 2013b), and multiple genes for H_2_-producing hydrogenases (Hyf; EC 1.12.1.4) (Trchounian and Trchounian, 2013), but lacks most genes for heme biosynthesis. Moreover, citrate synthase, aconitase, and isocitrate dehydrogenase genes in the TCA cycle are missing from the *Bac* genome (**Figure 5B**), disrupting glutamate biosynthesis. Therefore, these data, along with the specific malate requirement and unexpected glutamate export observed in strain PER-K23 cultures, portend a tightly entangled syntrophy between malate-consuming, glutamate-producing *Dhb* and the malate-producing, glutamate-auxotrophic *Bac*.

**Figure 5.**
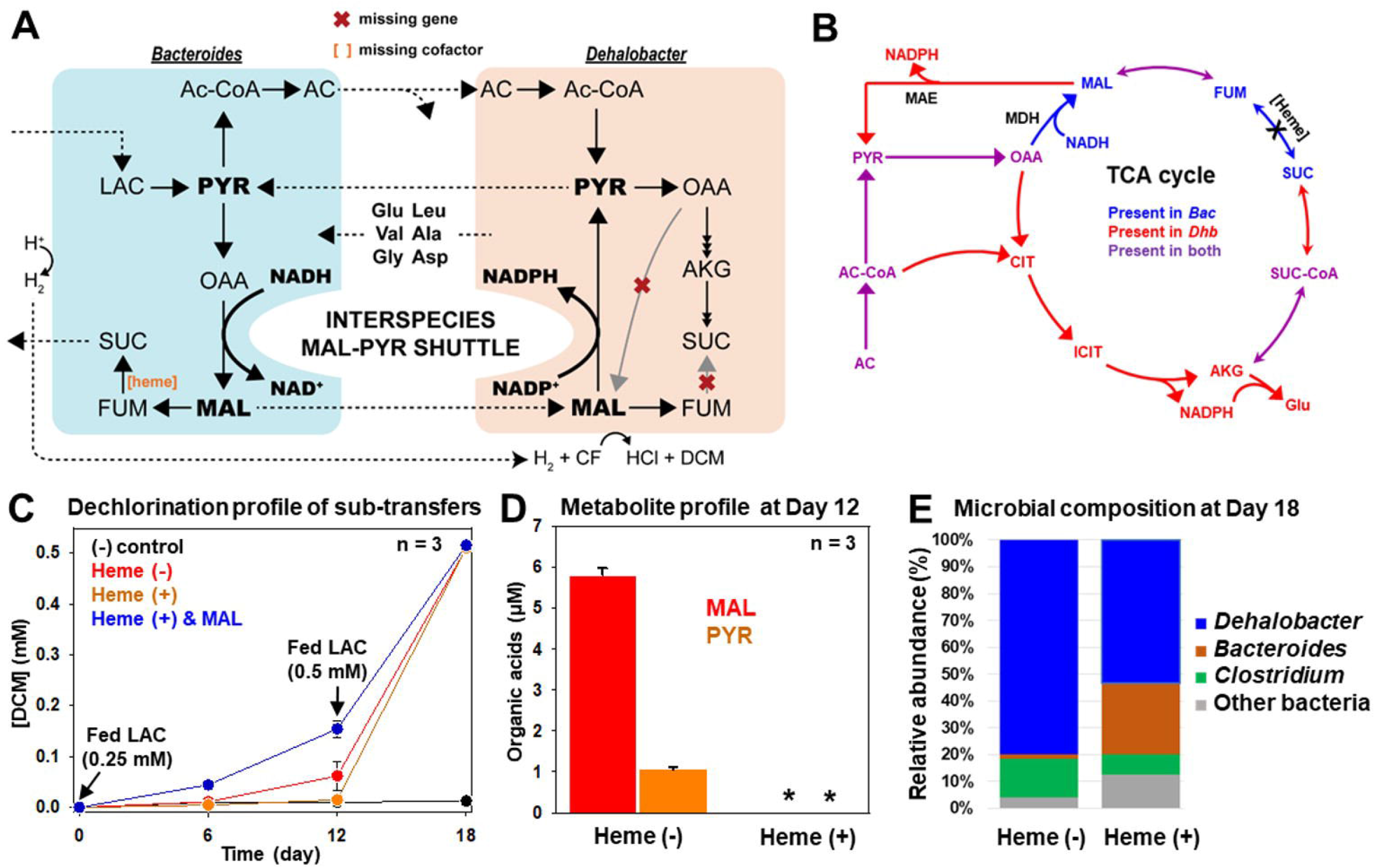
Proposed interspecies malate-pyruvate shuttle between *Dehalobacter restrictus* and the syntrophic *Bacteroides* sp. **(A)** Proposed syntrophy between *Dehalobacter restrictus* (*Dhb*) and the *Bacteroides* sp. (*Bac*) including syntrophic lactate fermentation/chloroform dechlorination and cross-feedings of malate, pyruvate, and amino acids. In the simplified co-culture model, lactate (LAC) is used as the sole electron donor and carbon source, and chloroform (CF) is used as the final electron acceptor. The red crosses (X) represent the missing genes in *Dhb* genomes; the orange square bracket [heme] represents the lack of heme cofactor in *Bac* due to the missing heme biosynthesis pathway. **(B)** Proposed genome streamlining between NADPH-restricted *Dhb* and glutamate-auxotrophic *Bac* demonstrated by complementary genes in the TCA cycle and in malate metabolism (present in *Dhb*, red; present in *Bac*, blue; present in both, purple). **(C)** Dechlorination profile of the 4% ACT-3 sub-transfer cultures added with only lactate (Heme(−)); lactate and heme (Heme(+)); lactate, heme, and malate (0.25 mM; Heme (+) & MAL). A second feed of lactate (0.5 mM) was given to all the sub-transfer cultures at Day 12 after supernatant sampling. **(D)** LC-MS analysis of malate (red bar) and pyruvate (orange bar) in the supernatants of Heme (−) and Heme (+) cultures at Day 12. The black asterisks represent that the metabolites are undetectable. **(E)** Microbial community composition (%) in Heme (−) and Heme (+) cultures at Day 18 determined using 16S rRNA gene amplicon sequencing. Abbreviations: AC, acetate; Ac-CoA, acetyl-CoA; DCM, dichloromethane; FUM, fumarate; OAA, oxaloacetate; SUC, succinate; MAE, malic enzyme; MDH, malate dehydrogenase. Data are means ± SE of three replicates in each experiment as shown on figures.

Based on literature and integrated multi-omics data, we proposed a metabolic interdependency between strain CF and *Bac* (**Figure 5A**). Given that (a) fermentative malate production from lactate is thermodynamically unfavorable under standard conditions (**Table 1**) and (b) heme is insoluble and never added to the growth medium, lactate is mainly fermented to H_2_, acetate and CO_2_ by *Bac*. Nevertheless, fermentative malate production from lactate would become thermodynamically favorable when (a) the H_2_ production is coupled to organohalide respiration by *Dhb* (**Table 1**) and (b) an efficient consumer is present to lower malate concentration. Therefore, *Bac* exports malate to consume excessive NADH generated from lactate and to facilitate NADPH regeneration in *Dhb*. In return, *Dhb* exports glutamate and other amino acids to consume excessive NADPH and to facilitate *Bac* anabolism. Since pyruvate was not accumulated in ACT-3 (**Figure S3**), the exported pyruvate from *Dhb* is likely recycled by other community members, including *Bac*, which shapes an intercellular metabolic cycle, enabling *Dhb* and *Bac* to indirectly trade NADH/NADPH across the membrane, resembling the function of a transhydrogenase (**Table 1**) (Voordouw *et al*., 1983).

### Decoupling interspecies malate-pyruvate exchange

We sought to validate the proposed malate-pyruvate exchange between *Dhb* and *Bac*. Given that *Bac* ferments lactate to malate as a result of heme-auxotrophy, heme addition to ACT-3 would enable further malate fermentation to succinate, disrupting the malate-pyruvate exchange and reducing available electron donors (i.e. H_2_) for strain CF (**Figure 5A**). Therefore, we monitored the effect of heme addition to ACT-3 sub-transfer cultures under limiting electron donor conditions (1-fold theoretical electron equivalent). After 12 days of incubation, chloroform dechlorination (50 μM) only occurred in ACT-3 sub-transfer cultures without heme addition (Heme (−) cultures) but not in cultures supplemented with 1 mg L^−1^ heme (Heme (+) cultures) (**Figure 5C**). Moreover, malate and pyruvate were only detected in supernatants of Heme (−) cultures but not in supernatants of Heme (+) cultures at Day 12 (**Figure 5D**). By contrast, Heme (+) culture supplemented with malate (0.25 mM) revealed significant chloroform dechlorination (0.2 mM). These data indicate that dechlorination in Heme (+) cultures is limited by the shortage of electron donors and malate. Consistently, when we provided excessive electron donors (3-fold theoretical electron equivalent) to the cultures at Day 12, all three cultures depleted chloroform within 6 days. Finally, since fumarate reduction to succinate in *Bac* is coupled to proton motive force, heme supplementation will enable *Bac* to synthesize more ATP via efficient oxidative phosphorylation rather than via substrate-level phosphorylation (Macy *et al*., 1975, Madej *et al*., 2006) (**Table 1**), thereby reaching a higher cell density. Consistently, the relative abundance of *Bac* in Heme (+) cultures at Day 18 is 10 times higher than that in Heme (−) cultures (**Figure 5E**). Therefore, our data support the occurrence of malate-pyruvate exchange between *Bac* and strain CF.

## Implications for microbial ecology

In this study, the proposed malate-pyruvate exchange between *Bac* and *Dhb* resembles the mitochondrial malate-pyruvate shuttle in eukaryotic cells (Liu *et al*., 2002, MacDonald, 1995). In the malate-pyruvate shuttle, cytoplasmic pyruvate is first transported to mitochondria, and is reduced to malate via NADH-dependent malate dehydrogenase in the TCA cycle (**Figure 3A**). Malate is then exported to cytoplasm via an antiporter, and decarboxylated to pyruvate by MAE for NADPH regeneration. Although the paring of metabolic partners in nature can be random, the complementary gaps in the TCA cycle and malate metabolism between *Dhb* and *Bac* genomes (**Figure 5B**) are likely a consequence of genome streamlining driven by co-adaption. This claim is supported by the presence of malate dehydrogenase and succinate dehydrogenase/fumarate reductase genes in most *Peptococcaceae* but not in *Dhb* (dashed boxes in **Figure 1**). Also, the oxidative TCA cycle is present in the genomes of all available *Bacteroides* isolates but not in *Bac*. Accordingly, this study has found a relevant example in biology, the interspecies malate-pyruvate shuttle, to support an endosymbiotic hypothesis in which mitochondria originated from a mutualistic interaction mediated by organic acid exchange (Searcy, 2003). In conclusion, the data present in this study demonstrate that metabolic complementarity and redox constraints (i.e. NADPH regeneration) are important driving forces for amino acid exchange in anaerobic microbial communities. Furthermore, the successful model prediction justified our previous argument that the accuracy of metabolic annotation can be greatly improved through consideration of cofactor availability and enzyme promiscuity. Finally, finding *Dhb* as an amino acid producer in the parent consortium reinforces the caveat that observed physiology of isolate cultures can deviate from their physiology in natural niches. Therefore, integration of laboratory experiments with computational metabolic modeling offers great opportunities to decipher the metabolic interdependency of fastidious, or currently unculturable, microorganisms.

## Acknowledgements

Support was provided by the Government of Ontario through Genome Ontario SPARK Research Grant. We also acknowledge the BioZone Mass Spectrometry facility for UPLC-ESI-HRMS analyses. We are grateful to the gift of active *Dehalobacter restrictus* strain PER-K23 culture from Dr. Jun Yan at the Löffler Lab in University of Tennessee (Knoxville, USA).

## Author contributions

E.A.E, R.M, and P.H.W conceptualized this study. C.H. and K.C. reconstructed the *Dehalobacter* model. K.C. performed the Flux Balance Analysis. P.H.W. performed the experiments. K.N. performed the organic acid analysis. R.F. performed the LC-MS analysis. P.H.W. and N.V. proposed the mechanism of interspecies malate-pyruvate shuttle. E.A.E, R.M, and P.H.W wrote this paper with helps from all the authors. All the authors were participated in data analysis and discussion.

Supplementary information is available at ISMEJ’s website

